# Dissociable roles of the auditory midbrain and cortex in processing the statistical features of natural sound textures

**DOI:** 10.1101/2023.04.12.536666

**Authors:** Fei Peng, Nicol S. Harper, Ambika Prasad Mishra, Ryszard Auksztulewicz, Jan W.H. Schnupp

## Abstract

It is well established that sound texture perception takes advantage of a hierarchy of time-averaged statistical features of acoustic stimuli, but much remains unclear about how these statistical features are processed in auditory subcortical and cortical regions. Here, we compared the neural representation of sound textures in the inferior colliculus (IC), and primary and non-primary auditory cortex (AC) of anesthetized rats with texture morph stimuli which gradually add statistical features of increasingly higher order. We generated texture morphs for a representative subset of 13 sound textures, chosen to span the three principal component dimensions of a corpus of 200 natural texture recordings. For each of the 13 texture types, six different exemplar morphs were synthesized using different random seeds. All exemplars of each texture type have the same long term statistics, but they differ in their acoustic waveforms. Meanwhile, different texture types differ in statistical features as well as waveforms. An analysis of transient and ongoing multi-unit responses to our stimulus set showed that the IC units were sensitive to every type of statistical feature, albeit to a varying extent, but only a small proportion of AC units were overtly sensitive to any statistical features. Differences in texture types explained more of the variance of IC neural responses than did differences in exemplars, indicating a degree of “texture-type tuning” in the IC, but this was, perhaps surprisingly, not the case for AC responses. We also evaluated the accuracy of texture type classification from single-trial population activity and found that IC responses became more informative as more summary statistics were included in the texture morphs, while for AC population responses classification performance remained constant, and consistently lower, than for the IC. These results argue against the idea that AC plays an important role in encoding statistical features of natural sound textures.

## Introduction

How neural activity represents natural sounds at different stages of the auditory pathway is a central question in auditory neuroscience. At the lower stations of the auditory pathway, neural responses mostly reflect the physical properties of incoming sounds, such as spectral composition, modulation or binaural properties. However, it is generally assumed that at higher auditory stations, neural representations may become increasingly abstract, perhaps coding for classes of objects that are identifiable by the types of sounds they make, such as vocalizations of a particular type of animal (Grace et al., 2003; Montes-Lourido et al., 2021), rather than merely implementing a set of simple acoustic filters and feature detectors. For instance, neural responses in the primary auditory cortex (A1) carry less information about the spectro-temporal structure and sound identity of bird vocalizations than the IC neural responses (Chechik et al., 2006; Chechik & Nelken, 2012). IC neurons convey similar amounts of information about spectro-temporal patterns as they encode about sound identity, but in A1, the amount of sound identity information encoded by neural responses is thought to exceed the amount of spectro-temporal pattern information (Chechik & Nelken, 2012). Thus, the nature of the representation changes in ways that are still only poorly understood.

Sound textures have become a useful class of natural sounds to study these phenomena, because although they are of considerably higher complexity than standard laboratory sound stimuli, they are nevertheless fully described by a set of statistical features relating to the sound power spectrum (e.g., its mean, variance, skewness, kurtosis, etc.) which can be assumed to be constant over time. This makes it possible to capture a lot of the diversity of natural sound recordings with relatively modest sets of statistical feature coefficients (Mishra et al., 2020), or to generate synthetic texture sounds that mimic recognizable classes of natural sound textures with considerable fidelity (McDermott & Simoncelli, 2011). Statistical features of sound textures are therefore a very useful tool for studying the sensitivity of neurons along the auditory pathway to complex, naturalistic sounds in a systematic and well-parameterized manner. Indeed, the usefulness of statistical parameters for characterizing different texture classes may be due to the fact that neurons at higher stations of the auditory pathway may themselves be tuned to such statistical parameters.

The idea that various stations of the auditory pathway may exhibit certain forms of tuning for the statistical properties of sound is not new. Several previous studies have investigated statistical feature sensitivity, particularly in subcortical structures such as the inferior colliculus. For example, it has been shown that subcortical neurons can be sensitive to stimulus variance (Kvale & Schreiner, 2004; Lohse et al., 2020), modulation power spectra (Hsu et al., 2004), or the spectro-temporal correlation structure (Escabi et al., 2003, Sadeghi et al., 2019). Sensitivity to statistical features of textures can also be represented by neural population codes, such as neural ensemble correlation (Zhai et al. 2020). Similarly, a recent study from our group (Mishra et al., 2021) showed that sensitivity to all types of statistical texture features described by McDermont and Simoncelli (2011) can be observed in the transient and sustained multiunit activity of IC neurons, although to a varying extent depending on the feature.

Less is known about the neural sensitivity of cortical neurons to statistical features of sound textures. Evoked activity in auditory cortex measured with fMRI in humans (Norman-Haignere & McDermott, 2018) or by ultrasound imaging in ferrets (Landemard et al., 2020) in response to synthetic textures with a full set of statistical features closely resembles that evoked by corresponding natural sound texture recordings. However, the sensitivity of single or multi-units in the auditory cortex to different types of statistical features has not yet been described. Neurons in the A1 are certainly known to be sensitive to some types of statistical features of synthetic sounds, such as spectral contrast (Rabinowitz et al., 2011; So et al., 2020) or temporal correlations (Natan et al., 2017) of dynamic random chord sequences. The majority of individual neurons in A1 were selective for a subset of stimuli with specific spectro-temporal density and cyclo-temporal statistics, corresponding to a range of water-like percepts (Blackwell et al., 2016). However, the neural population responses were stable over very large changes in stimulus statistics (Blackwell et al., 2016). Cortical neurons also prefer envelopes with a more natural 1/f envelope distribution to unnaturally slow or unnaturally fast modulations (Garcia-Lazaro et al., 2006). Since synthetic textures become increasingly similar to natural sound textures as more higher-order statistical features are incorporated in the synthesis, one might therefore similarly expect AC neurons to prefer synthetic textures with many, appropriately chosen high-order statistical features over simpler synthetic variants with fewer statistical features.

The current study builds on our previous work characterizing statistical feature sensitivity in the auditory midbrain (Mishra, Peng, et al., 2021), and aims to investigate how the neural sensitivity to different types of statistical features of sound textures changes along the auditory pathway, and how this feature sensitivity may contribute to the encoding of sound texture identity in different regions. We tested two key hypotheses: (1) neural responses in both IC and AC might become better at distinguishing sound texture types as the number of statistical features available increases, and (2) AC, being higher up in the processing hierarchy of the auditory pathway, might be better able than IC to generalize across different exemplars of the same texture type when categorizing texture stimuli. To pursue these questions, we selected 13 sound texture types so as to sample fairly evenly from the principal component space of sound textures that we had previously described (Mishra, Harper, et al., 2021). For each texture type, we systematically generated synthetic variants in a step-wise way, gradually incorporating statistical parameters of increasingly higher order. Multiple different exemplars were synthesized for each type, starting from different random seed values. We then exposed anesthetized rats to the synthetic textures while recording the electrophysiological multiunit activity in the IC, A1, and non-primary (non-A1) auditory cortex. We analyzed whether individual multiunits were sensitive to the changes of statistical features, and examined the selectivity of multiunits with respect to texture types and/or exemplars. We also tested whether the texture type classification performance based on single-trial neural responses changed over the increasingly complex synthetic variants, and compared the classification performance based on activity in the IC, A1, and non-A1.

## Materials and Methods

### Sound stimuli

The procedures for selecting the natural sound textures and generating the stimuli were described in our previous study of the IC (Mishra et al., 2021). Briefly, we selected 13 representative texture sounds, evenly sampling the space of 3 statistics inferred from a database of over 200 natural textures. The selected natural textures had the following descriptors: “applause”, “barn swallow calls”, “cackling geese”, “church bells”, “burning wood sticks”, “fireworks”, “foot-steps in water”, “frogs at night”, “galloping horses”, “stirring liquid in a glass”, “lawnmower”, “xylophone”, and “tin can”. Following the sound synthesis procedures described by McDermott & Simoncelli (2011) to generate synthetic variants, Gaussian white noise was gradually morphed to increasingly resemble the natural texture by sequentially incorporating the following sets of statistical features: the sound power spectrum (Power), variance (+Var), skew and kurtosis (+S.K), cochlear correlation (+Coch.Corr), modulation power (+Mod.power), and modulation correlation (+C1+C2). Segments of 1.5 s duration of each increasingly complex synthetic variant were concatenated with a 10 ms cosine cross fade. Below we refer to these timepoints when statistical features change as “transitions”. A segment of 1.5 s duration, randomly selected from the original sound (Ori), was appended at the end. We refer to one stimulus which transitions in this way from Gaussian noise to the full, natural texture sound as a “texture morph”. The duration of each of these morphs, including all the synthetic variants and the Ori, was 10.5 s. The stimuli were normalized to have the same RMS power in each of the synthetic variants, and delivered at 80 dB SPL. Using different random seeds, we generated 6 exemplars for each synthetic variant for each texture, resulting in a total of 78 different morphs (13 textures x 6 exemplars). During the recording experiment, each morph was presented 10 times, for a total of 780 stimuli presented in a random order.

In addition to recording responses to the texture morphs, pure tones (duration 100 ms with 5 ms rise/fall ramps) with frequencies ranging from 0.5 to 32 kHz in 1/4 octave steps, and sound levels varying between 30 and 80 dB SPL in 10 dB steps, were presented to estimate the best frequency (BF) of each unit. Each tone was played 10 times.

### Neurophysiological recording

Young adult female Wistar rats (N = 10, body weight 250-280 g) were used in the study. Experimental procedures and protocols were approved by the City University Animal Research Ethics Sub-Committee and conducted under license by the Department of Health of Hong Kong [Ref. No. (19-31) in DH/SHS/8/2/5 Pt.5].

Extracellular recordings were collected from the AC from 5 rats. For the purposes of systematic comparison, we also present here IC recordings from a further 5 rats which were collected during a previous study (Mishra et al., 2021) using identical stimuli and identical techniques, except for the type and placement of extracellular recording electrodes. Animals were anesthetized by an initial injection of ketamine (80 mg/kg) and xylazine (12 mg/kg), and the anesthesia status was maintained by a continuous injection of the mixture of 0.9% saline solution of ketamine (17.8 mg/kg/h) and xylazine (2.7 mg/kg/h) with a syringe pump at a rate of 2.1 ml/h. Body temperature was kept at 38 °C using a heating pad (RWD Life Science, Shenzhen, China). The animals were head-fixed with hollow ear bars in a stereotactic frame, and placed inside a sound-proof chamber. Normal hearing sensitivity was confirmed based on the threshold of the auditory brainstem responses to clicks presented broadband at a rate of 23 Hz (< 30 dB SPL).

For the IC recordings (Mishra et al., 2021), a craniotomy had been performed just anterior to lambda, and extracellular activity had been recorded using a single-shank 32-channel silicon electrode with 50 μm spacings between recording sites (ATLAS Neuroengineering, E32-50-S1-L6) inserted into the IC in a dorsal-ventral direction through the overlying cortex. For the AC recordings performed for this study, a rectangular 5 × 4 mm craniotomy was performed to extend from 2.5 to 7.5 mm posterior from bregma and with a medial edge 2.5 mm from the midline to expose the AC (Polley et al., 2007). AC neural responses were collected using 64-channel arrays, composed of four shanks each with 16 electrodes, with 100 μm spacing between electrodes (ATLAS Neuroengineering, E64-100-S4-L6-600). Both types of neural raw signals were amplified using a PZ5 preamplifier and recorded at a sampling rate of 24414 Hz with an RZ2 system (Tucker-Davis Technologies).

### Data analysis

#### Preprocessing of the AMUA

Analog multiunit activity (AMUA) was used as a measure of neural activity in the frequency range occupied by spikes (Schnupp et al., 2015). To obtain AMUA, the raw signal was bandpass filtered between 300 and 6000 Hz by a third-order zero-phase Butterworth filter. The absolute value of the filtered signal was taken, and the absolute signal was downsampled to 2 kHz with appropriate low-pass filtering. This resultant raw AMUA measure quantifies the electrode signal amplitude in the 300-6000 Hz band which is normally occupied by action potentials. To decide whether the AMUA at a given recording site was responsive to acoustic stimulation, we compared the onset response (the mean AMUA amplitude within the first 100 ms following stimulus onset) with the pre-stimulus baseline of spontaneous activity (quantified by as the mean AMUA amplitude over the 500 ms of silence preceding stimulus onset). For each of the 13 textures, we pooled the responses across all 10 trials for all 6 exemplars, and we determined whether the AMUA onset response (up to 100 ms after the sound onset) was significantly different from the baseline (500 ms before the sound onset) AMUA amplitude using a paired sign rank test. The 13 p-values obtained for each of the textures were combined into a single p-value using Fisher’s method, and multiunits were deemed responsive if the p value reached the highly conservative threshold of p < 0.0001. There were 480 IC multiunits (out of n = 480) and 310 AC units (out of n = 576) which showed significant onset responses, and these were analyzed further in this study. As a final preprocessing step, we computed the “response strength” of each multiunit as the AMUA amplitude, normalized relative to baseline activity by subtracting the mean of the pre-stimulus baseline and dividing the result by the standard deviation of the pre-stimulus baseline over time. This response strength measure is therefore effectively a z-score of AMUA amplitudes centered on the baseline activity.

#### Physiological classification of AC multiunits

To classify the recorded AC units into putative A1 or non-A1 units, we used stereotaxic recording positions in combination with neural response properties observed during frequency response area (FRA mapping with pure tones, including, response peak amplitude, onset latency, and response duration (Polley et al., 2007). BF was defined as the frequency which evoked the maximum AMUA at a sound level of 80 dB SPL. Response peak amplitude was defined as the peak amplitude of the AMUA response in a time window up to 200 ms after tone onset averaged over all tone stimuli. Onset latency was defined as the point in time where the rising slope of the AMUA post stimulus time histogram, averaged over all frequencies and binned at 0.5 ms, crosses the level of the mean evoked response. Response duration was defined as the time difference between the onset latency and the time point at which the excitatory response had decreased back to below the mean evoked response. We compared these features across the units, and identified tonotopic gradients to confirm the location of A1. In the rat, the tonotopic gradient runs from low frequencies rostrally to high frequencies caudally (Polley et al., 2007). Taking recording position and all the above features into account, we classified 209 units as A1 and 101 units as non-A1. The mean (± standard error) response strength peak amplitude in A1 was 0.55 (± 0.04), compared to 0.21 (± 0.02) for non-A1. The mean onset latency was shorter in A1 (21 ± 0.5 ms) than non-A1 (23 ± 1 ms), and the response duration was also shorter in A1 (41 ± 1 ms) than non-A1 (53 ± 2 ms).

#### Neural sensitivity to the changes of statistical features

To ask whether the neural responses are sensitive to these statistical features, we hypothesized that such a sensitivity should manifest in changes in neural activity following changes in statistical features of the texture morph. We used a bootstrapping method which we described in our previous study of IC neural responses (Mishra et al., 2021) to examine whether any observed response amplitude changes were statistically significant. We looked for both transient (0 to 50 ms after the transition) and sustained (0.5 to 1.5 s after the transition) changes in firing accompanying changes in statistical parameters. Briefly, for the transient change, we averaged the measured response strength values (quantified as described above) over each of the 10 repeats of each exemplar and over each of the 6 exemplars of each texture, and computed the absolute difference (50 ms before and after the transition) of the averaged responses to compute the “observed transition response”.

To judge the statistical significance of observed changes we compared them against an estimated null distribution, which was computed by choosing simulated “null” transition times picked at random, uniformly, from 950 ms to 150 ms before the given transition. For each simulated transition time we computed response changes in an identical manner as for the actual transition time. This computation was repeated 1000 times to generate a distribution of simulated null transition responses, and the p-value for the “true”, observed transition response was computed as the percentile of the true transition response value in the distribution of null transition values. To be deemed significantly sensitive to a particular change in statistical parameters, a multiunit had to generate p-values < 0.05 for at least 4 of the 13 textures. To confirm the robustness of this “N out of 13” criterion, we performed a “sanity check” by investigating how many false positives we would generate if the actual p-values were replaced by simulated null-distribution p-values drawn at random form the values generated during the bootstrap test. “False transition responses” compared the response 50 ms before the transition and the response 100 ms to 50 ms before the transition. We conducted these this test on all 6 transitions and all 480 IC multiunits, 209 A1 multiunits and 101 non-A1 multiunits in our sample, and we obtained 1 false positive result on 1 transition for 1 multiunit in the IC, 4 false-positive results on 3 transitions for 3 multiunits in the A1, and 1 false-positive result on a single transition for a single multiunit in the non-A1. These low false positive rates confirm the high specificity of our analysis.

To quantify the strength of ongoing, “steady-state” responses, we averaged the AMUA strength in a 1s wide time window preceding the next transition. These values were then compared using a bootstrap test. For each texture, we resampled the averaged AMUA from the 60 samples for each texture (10 trials for each of the 6 exemplars) with replacement, and then calculated the mean of these 60 numbers. Repeating this process 1000 times, we obtained a bootstrap distribution of the mean ongoing AMUA amplitudes. Ongoing responses before and after a stimulus transition were deemed statistically significantly different if the pre- and post-transition bootstrap distributions overlapped by less than 5%. As with the transient responses, a multiunit needed to exhibit significant differences in the pre- and post-transition ongoing responses for at least 4 of the 13 textures for the multiunit to be considered sensitive to a given type of statistical features.

#### Neural selectivity for texture types over exemplars (analysis the variance of neural response)

Different exemplars of the same texture type exhibit the same long term statistical features but have different waveforms. If an individual multiunit could be described as sensitive to statistical features that distinguish some texture types from others, one would expect that this feature selectivity should show a degree of invariance with respect to exemplars, that is, that multiunit’s responses to different exemplars of the same texture should be more similar than its responses to different textures. This intuition is amenable to an analysis which computes and compares the proportion of the variance in neural responses that can be “explained” by changes either in stimulus texture type or by changes in exemplar while holding the texture type constant. As was pointed out in a previous study of neural responses to synthesized, naturalistic visual textures in the visual cortex (Ziemba et al. 2016), the appropriate framework for this purpose is that as of a nested ANOVA, as each of the texture exemplars is a member of only one type of textures, so that trials during which any one exemplar was presented are “nested” within textures.

We therefore computed the percentage variance of neural responses across texture types and across exemplars within a texture type of each unit. The neural responses were averaged for each trial in the ongoing time window (100 to 1500 ms after the feature transition). Total variance indicates the variance of neural responses to all stimuli (13 texture types x 6 exemplars x 10 trials); the variance across texture types indicates the variance of neural responses around the average response across exemplars for each texture type and over trials; while the variance across texture exemplars indicates the variance of neural response around the mean response for each texture exemplar. The variance explained (VE) by texture type was computed as the ratio of variance across texture types and total variance, and the VE by exemplar was computed as the ratio of variance across texture exemplars and total variance. It is important to be aware that these VE estimates, much like mutual information estimates (Treves and Panzeri, 1995), can be subject to upward bias because the estimates can be inflated by the effects of sampling noise. We used a bootstrapping method to bias correct our VE estimates and to determine whether the observed VE estimates were significantly greater than zero. We randomly shuffled the type and exemplar labels which leads to an expected VE of zero, and re-computed the observed VE. This procedure was repeated for 1000 iterations to estimate the null distribution for the VE estimate. The mean of this null distribution was subtracted from the original VE to calculate the bias corrected VE value, and the VE was deemed significantly greater than zero if the original VE estimate exceeded the 95th percentile of the bootstrapped null distribution.

#### Classification decoding of single-trial neural ensemble responses

To investigate whether the time-averaged single-trial neural responses of groups of units could be used to classify stimuli across the 13 texture types, we built a simple Gaussian decoder to quantify the neural classification performance. The decoding methods were adapted from previous research on the classification of visual textures based on neural activity in the visual cortex (Ziemba et al., 2016). We trained the decoder using the neural response to the original texture, and then tested the decoder on the time-averaged single-trial neural responses to the texture morphs in a 13-alternative forced-choice task, so the chance performance was 1/13. For the “training”, we computed the mean response strength of a neural population sample (1, 10, 30, or 80 units) over all exemplars and trials and over the full duration of 13 original sounds as the prior, which is analogous to establishing an internal fixed template for each texture type. Then, for the decoding, neural responses to each synthetic variant, single-trial neural responses were time-averaged over a randomly selected time window of steady state responses (50, 100, 300, 500 or 1000 ms). We estimated which of the 13 textures was most likely to have been generated by the response of a neural population to the original texture type, under the assumption of independent Gaussian variability around each of the estimated mean response strengths for different texture types. To quantify overall classification performance, we calculated the mean classification performance over all trials and textures, and repeated this process 1000 times for different, randomly selected subsets of 1, 10, 30, or 80 units. Our final classification performance estimate for each ensemble size was the mean classification performance of each of the 1000 simulated neural ensembles. The 2.5th and 97.5th percentile of the distribution of 1000 simulated ensembles served as confidence intervals for the mean classification performance estimate.

Because this performance was repeated separately with different response durations, we were also able to use the results from this analysis to estimate the “minimum classification time” as the shortest response time window for which the lower bound of the confidence interval of the mean classification performance exceeded the chance performance level.

## Results

Using the stimuli and methodology described in the Introduction and Methods sections, we recorded and analyzed the neural activity of 480 multiunits in the IC of 5 rats (which had previously been described in Mishra et al., (2021)), and 310 multiunits in auditory cortex of a further 5 rats. The cortical multiunits were classified into 209 A1 and 101 non-A1 multiunits using the criteria described in the Methods. Figure 1 shows representative examples of multiunit responses, one each taken from the IC, A1, and non-A1. For each multiunit we show responses to 6 exemplars of 4 different textures. The figure indicates that the responses of the multiunits lock to acoustic features in the texture morphs, that these time-locked responses may or may not coincide with the transition times when the statistical features of the sounds change, and that the amount of activity evoked by the stimuli can vary considerably between different textures, different exemplars of the same texture, as well as different morphs of the same exemplar.

**Figure 1.**
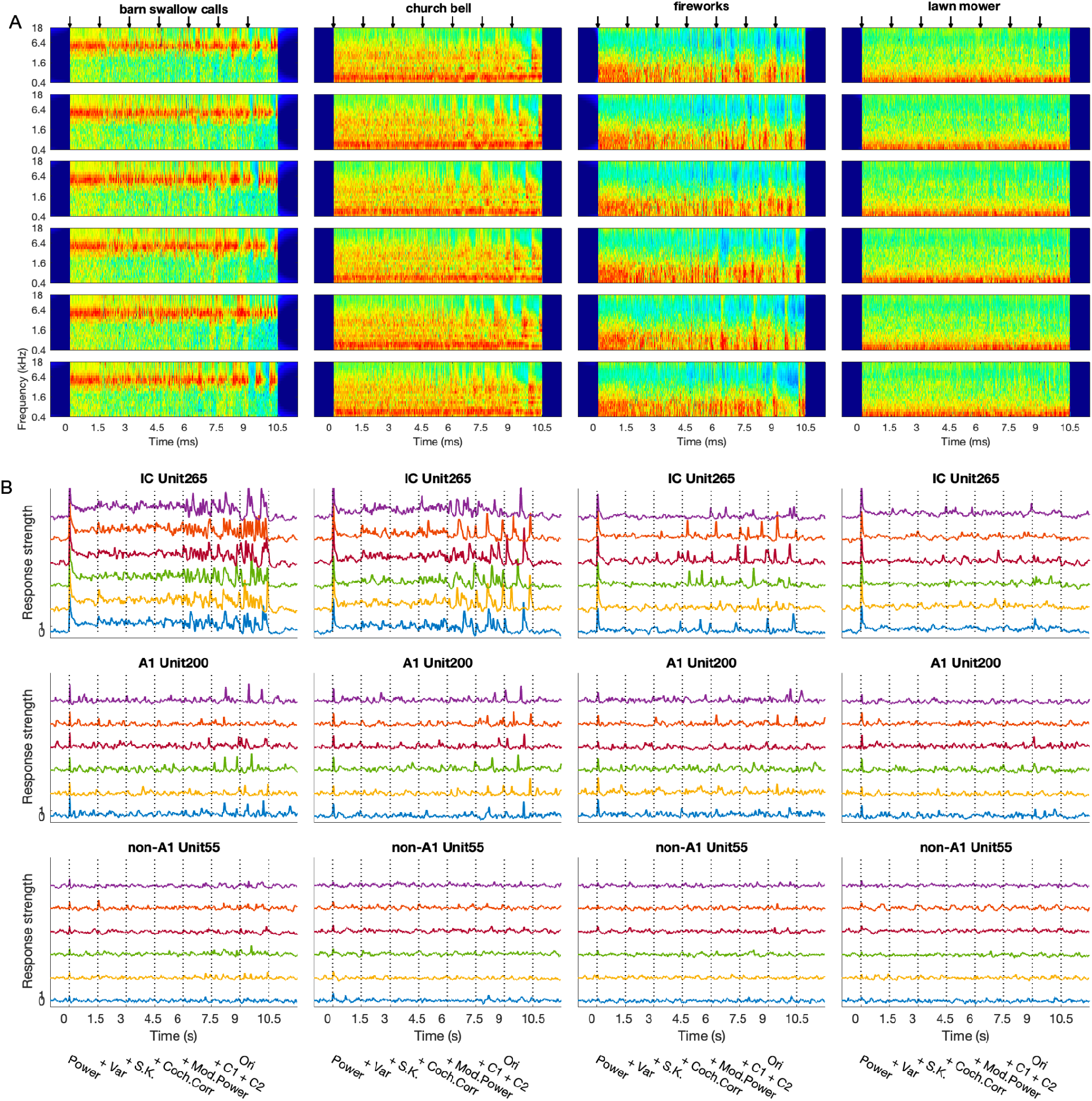
Cochleagrams of a subset of the stimuli alongside representative examples of neural responses. (A) Cochleagram for four of the 13 textures (“barn swallow calls”, “church bell”, “fireworks” and “lawn mower”). Each row represents a different exemplar. The arrows above indicate transition times when the statistical features of the texture morphs changed. (B) Normalized AMUA responses for three representative multiunits, one from IC, one from A1, and one from non-A1, to the four textures shown above. Each of the colored lines shows the mean response, averaged over 10 repeats, to a different exemplar. The dashed lines indicate the transition times when the statistical features of the texture morphs changed. From left to right, the texture morphs correspond to Power, +Var, +S.K., +Coch.Corr, +Mod.Power, +C1+C2, and Ori, respectively.

### Sensitivity of individual unit to the changes of statistical features of sound textures

First, we examined how sensitivity to each of the statistical feature types investigated was distributed among multiunits recorded at each of the anatomical stations recorded. The statistical features that characterize auditory textures are often described as forming a “hierarchy” (McDermott & Simoncelli, 2011), with, for example, correlations between envelopes of different frequency bands being in a sense a “higher order” feature than the variance or kurtosis of the envelope amplitudes in individual frequency bands. One might expect that sensitivity to features of increasingly higher order emerges only gradually along the ascending auditory pathway. However, our investigation into how common sensitivity to the different types of statistical features is among multiunits in midbrain and cortex respectively does not bear out this expectation, as can be seen in Figure 2. The histograms in Figure 2 show the proportion of multiunits which were deemed sensitive to changes in a given type of statistical feature of the sound textures in each of the stations of the auditory pathway examined. Feature sensitivity was determined according to either transient or sustained responses using the bootstrap test described in the methods section. Inspection of the figure reveals the following: Firstly, sensitivity to changes in statistical features is widespread in the IC, and manifests in both transient responses and sustained changes in firing rates. Even sensitivities to high-level features, including modulation correlations (+C1+C2), are not uncommon in the IC, being observed in up to 20% of multiunits examined. Secondly, and perhaps surprisingly, an overt sensitivity to changes in almost all types of statistical features is much less common in cortical responses than in IC responses, and this is even more pronounced in non-A1 than A1 multiunits. With the exception of the +Var feature, the proportion of cortical multiunits exhibiting a clear, overt sensitivity to changes in statistical texture features never exceeds a few percent. While the activity of cortical neurons is often clearly modulated by, or responsive to, the texture stimuli used in this experiment (consider the example of the A1 multiunit in Fig 1B), these cortical neurons only rarely respond directly to changes in statistical features.

**Figure 2.**
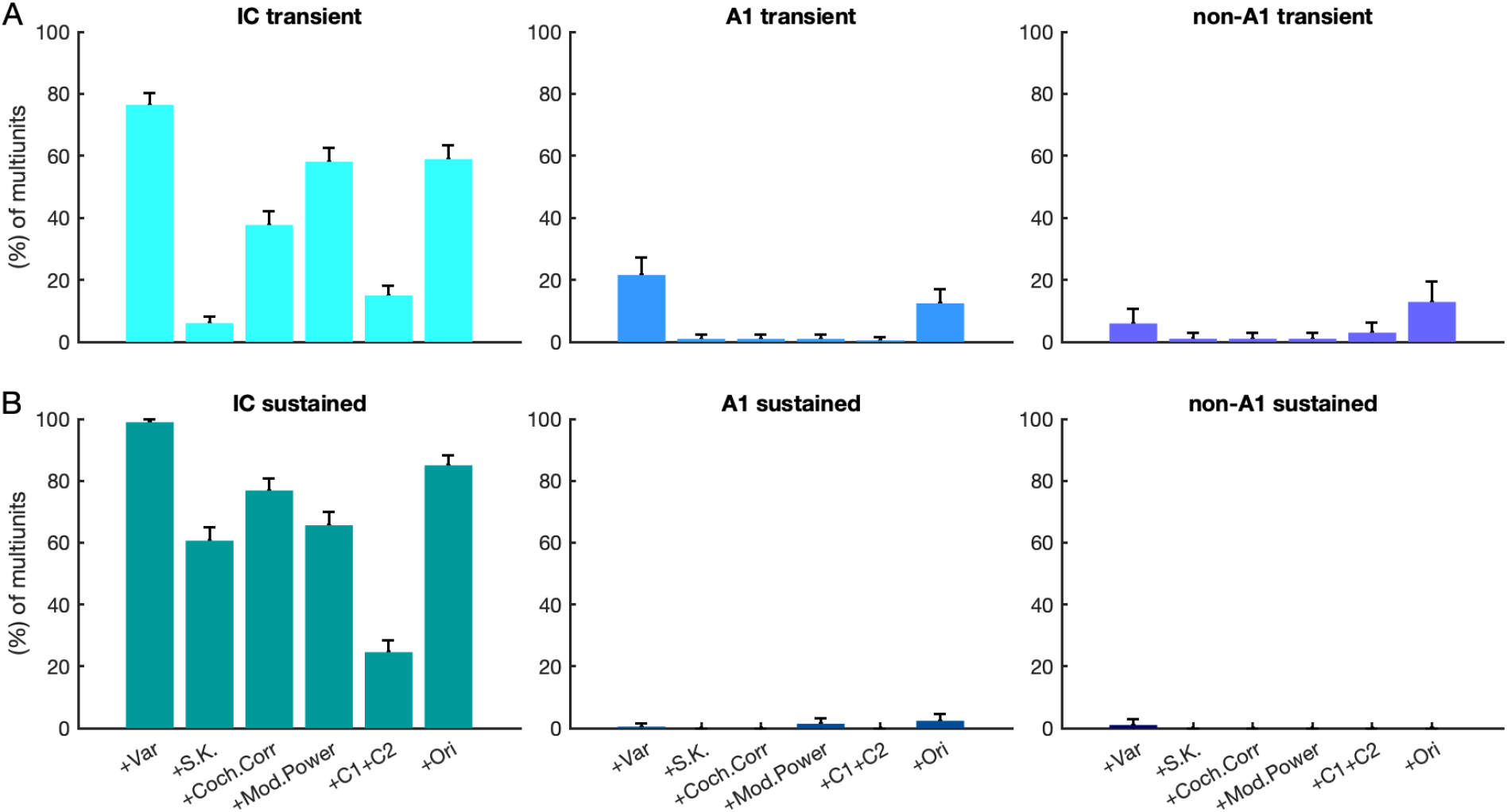
The percentage of multiunits showing significant changes in transient response (A) and sustained response (B) across texture morphs. Three columns represent the neural response in the IC, A1 and non-A1, respectively. Error bars represent 95% Wilson confidence intervals.

### IC units are more selective to texture types over exemplars than AC units

The analysis shown in Figure 2 tabulated the proportion of IC, A1 and non-A1 cortical multiunits that exhibited statistically significant changes in neural activity in response to changes in statistical features which were observed across multiple exemplars. It revealed that an overt sensitivity to changes in statistical texture features was widespread in the IC but rare in cortex. This suggests that cortical neurons may also be less able to discriminate between types of sound textures than IC neurons. To investigate this possibility, we performed an analysis of variance (nested ANOVA, see methods section) to quantify how much of the variance in neural responses can be attributed to (“explained” by) changing stimulus texture type or stimulus exemplar respectively. This analysis, carried out on the responses to the final part of the stimuli that contained the original (Ori) texture recordings, allowed us to compute the proportion of response variance explained by stimulus texture type and by stimulus exemplar within a texture type respectively, for each unit.

Figures 3A and B give an intuition for this analysis by showing example responses for two multiunits from IC, A1 and non-A1 respectively. The multiunits were chosen to have either high (95th centile; Figure 3A) or average (median; Figure 3B) VE by texture type. Each small black dot shows the response to a single stimulus presentation on the y-axis, and the responses are arranged by exemplar and texture type on the x-axis. First consider the IC multiunit shown in the leftmost panel of Figure 3A. For this multiunit, the response strengths for different texture types are clearly distinct. The variability from exemplar to exemplar within one texture type appears smaller than that from texture type to texture type, and the scatter of individual responses around the mean for each exemplar is similarly relatively small compared to the mean differences in responses for different texture types. It is therefore unsurprising that, for this particular multiunit, stimulus texture type accounted for (“explained”) a very large proportion (82.9%) of the variance in neural responses.

**Figure 3.**
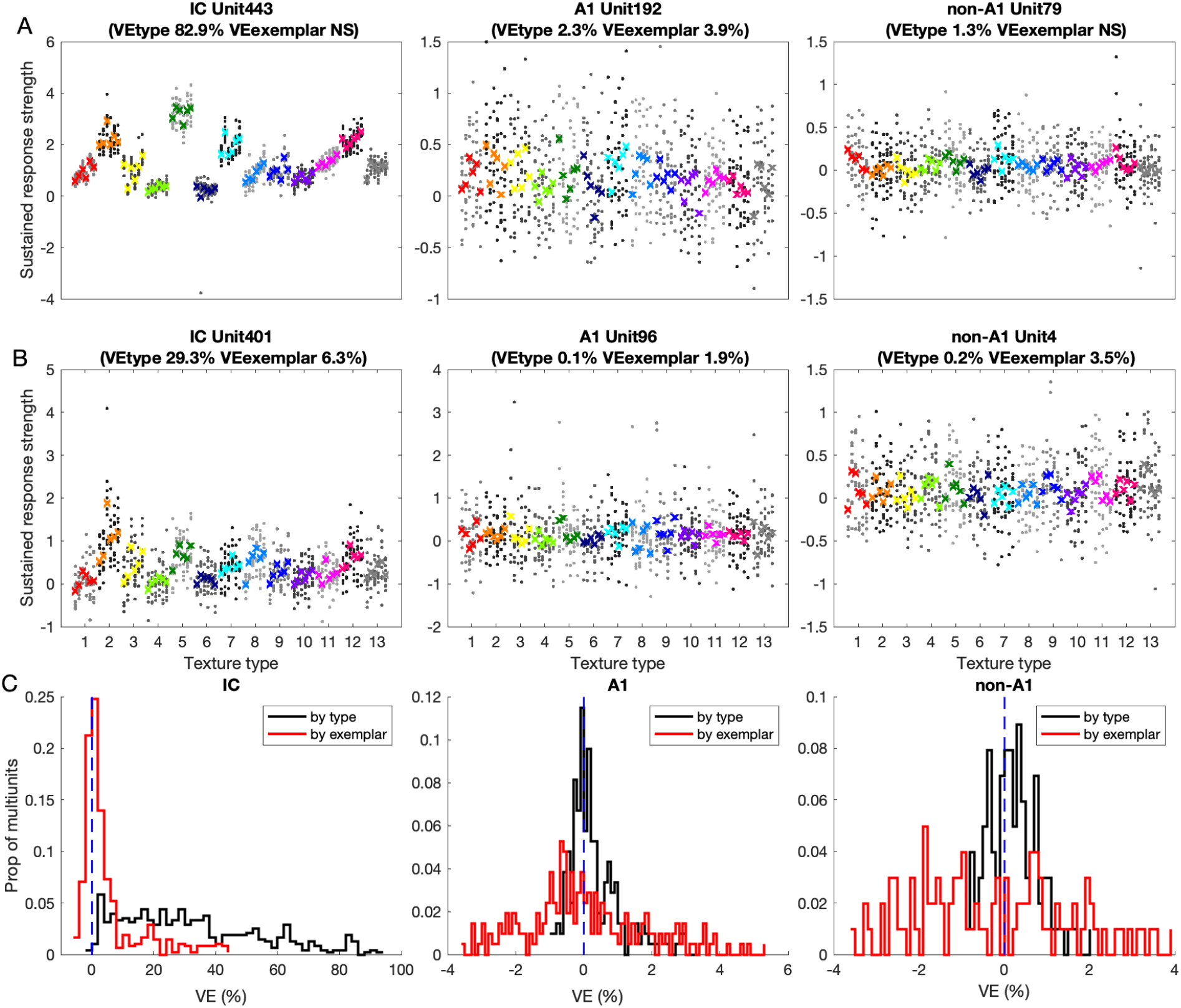
Variance analysis of individual multiunit responses to the original sounds. (A) Responses of one representative example unit from each region, chosen at the 95th centile of the distribution of bias VE values by texture type. Each dot indicates the averaged response strength in the sustained time window for one trial, the colored crosses indicate the mean response strength over trials for one exemplar. Each color represents the responses to one texture type indicated at the horizontal axis (with 13 texture types in total), and each column represents the response to one exemplar (with 6 exemplars per texture type). (B) Responses of one unit from each region, chosen to be an example of a strongly texture selective unit, at the 50th centile (median) of the VE by texture type. (C) Distribution of the VE by type (red line) and by exemplar (black line), respectively. The dashed blue line marks corrected VE = 0.

The IC multiunit shown in the leftmost panel of Fig 3B is chosen to be highly representative, with a VE by texture type that is the median of our IC dataset. While there is a lot more overlap in the distribution of responses to textures of different types, we nevertheless observe that some texture types (eg 2, 5 and 7) tend to evoke systematically stronger responses than others (eg 1, 4, 6, …). Consequently, observing the firing of this multiunit could clearly help discriminate stimulus texture types, and the VE by texture type of 29.3% reflects this. Note also that for both the examples of IC multiunits shown in Figure 3A and B, the VE by exemplar is noticeably smaller than the VE by texture type, which reflects the fact that response variability from exemplar to exemplar within one texture type tended to be smaller than the variability from one texture type to another. In that sense, these IC units appear to “generalize”: to give just one conspicuous example, the IC multiunit in Figure 3A fired consistently more strongly to any exemplar of texture 2 than to any exemplar of texture 4.

The examples of IC multiunits in Figure 3A and B thus show that the firing of IC units is often reasonably good and sometimes very good at discriminating texture types in a manner that can be said to exhibit a degree of invariance or generalization across texture exemplars. If we now contrast this with the examples for A1 and non-A1 multiunits shown in the middle and rightmost panels of Figure 3A and B, it is apparent that the ability of cortical neuron firing to discriminate texture types is remarkably poor when compared to that of the IC. For cortical units, the VE by texture type was invariably quite small, and the VE by exemplar was even smaller, indicating that these cortical units were more sensitive to texture types. As for the VE by texture type, we found that 99%, 14%, and 6% of the units in the IC, A1, and non-A1 were significantly (p<0.05) larger than the null bootstrapping distribution, however, for the VE by texture exemplar we found that only 39%, 11%, and 9% of the units in the IC, A1, and non-A1 were significantly larger than the null bootstrapping distribution. Accordingly, the VE by texture type was significantly (permutation test, p < 0.05) larger than those of VE by exemplars for units in the IC, A1, and non-A1, respectively (Figure 3C).

The variance analysis shown in Figure 3 suggests that IC units are substantially more sensitive to statistical features of textures than AC units. Note that the same recording methods and devices and the same inclusion criteria were used to identify sound responsive multiunits in the AC and the IC so the dramatically lower texture selectivity observed here for AC cannot be attributed to biases in the data acquisition. If sensitivity to statistical features drives this discrimination of texture types seen in IC multiunits responses, then these multiunits might become better able to discriminate texture types, as well as more invariant across exemplars, as the number of statistical features available increases. The VE by type might then increase, and the VE by exemplar might shrink, as the number of statistical features available to discriminate texture types increases. This possibility is testable by repeating the VE analysis just described on the responses to texture morphs of increasingly higher order.

In Figure 4, we plot, for each multiunit in the IC (Figure 4A), A1 (Figure 4B) or non-A1 (Figure 4C), VE by exemplar along the x-axis against VE by type along the y-axis, computed for the sustained responses to synthetic variants with features of increasingly higher order as shown above each feature panel. We use different colors depending on whether multiunits exhibited VE values that were significantly above zero for VE by type (black), VE by exemplar (red), both (blue) or neither (light gray). Multiunits for which the VE by type is larger than the VE by exemplar will lie above the main diagonal. We observe that, for the IC, somewhat in line with our prediction, the proportion of units for which VE by type exceeds VE by exemplar at each texture morphs increases from 83.5% for the Power-only morph to 88.5% in the +Var condition, but it then does not change much (+S.K.: 87.3%, +Coch.Corr: 89.4%, + Mod.Power: 87.7%, +C1+C2: 88.1%) until we reach the Original (Ori) condition where it jumps to 93.1%.

**Figure 4.**
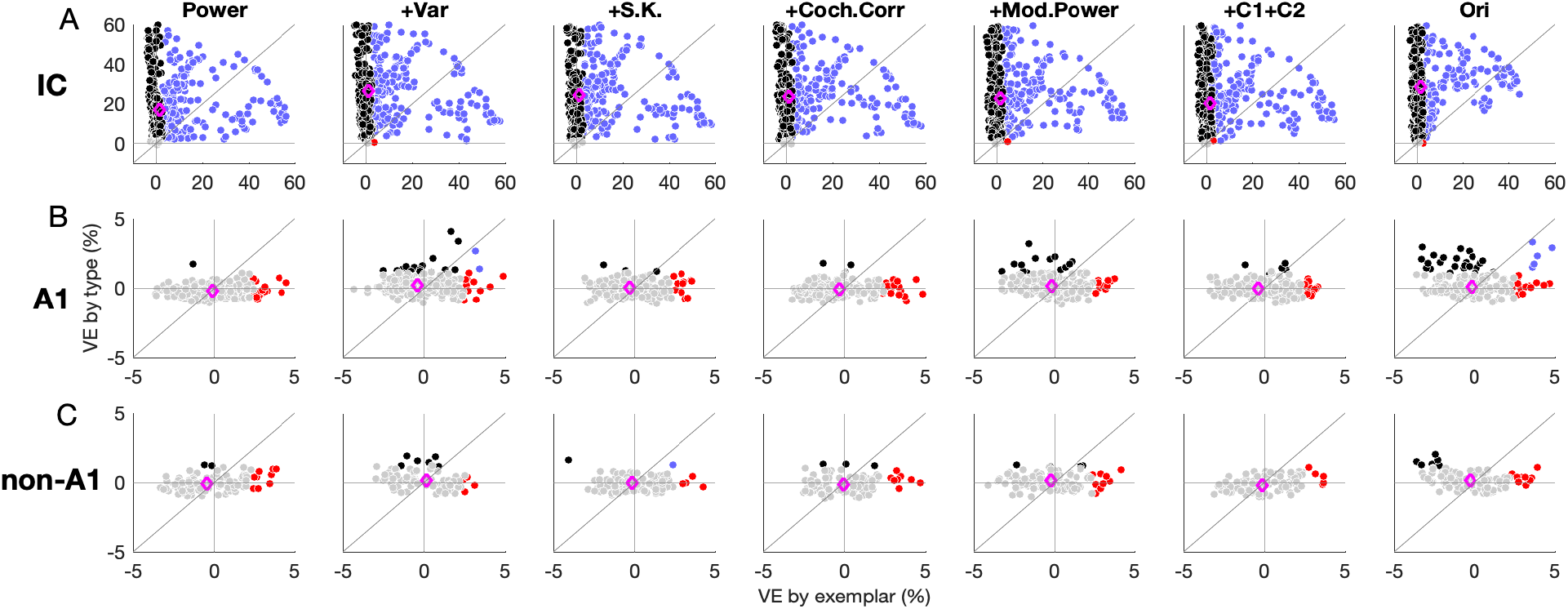
Percent VE by exemplar (x-axis) plotted against VE by texture type (y-axis) for IC (A), A1 (B) and non-A1 (C) units, for each type of texture morph. Each teal or blue dot shows the VE values for the sustained responses of one IC or cortical multiunit to the texture morph indicated above each panel respectively. The magenta diamond indicates the median VE value across units.

In the IC the proportions of multiunits units showing VE by texture type significantly above zero were 94.5% for Power, 99.2% for +Var, 98.3% for +S.K, 97.7% for +Coch.Corr, 98.5% for +Mod.Power, 99.0% for +C1+C2, and 99.2% for Ori. The VE type in the Power condition was significantly smaller than that at all the other stimulus conditions (permutation test, p < 0.05). In comparison, the proportions of multiunits exhibiting significant VE by exemplar were 42.3% for Power, 38.5% for +Var, 40.0% for +S.K, 37.1% for +Coch.Corr, 41.0% for +Mod.Power, 41.0% for +C1+C2, and 35.6% for Ori. The VE by exemplar in the Power condition was significantly larger than that at all the other stimulus conditions (permutation test, p < 0.05).

In the A1, the proportion of multiunits with significant VE by type or by exemplar were much smaller than in the IC. The observed proportions by type were 0.5% for Power, 0.4% for +Var, 0.5% for +S.K, 1% for +Coch.Corr, 4.3% for +Mod.Power, 1.4% for +C1+C2, and 12.4% for Ori. The proportions for significant VE by exemplar were 3.4% for Power, 3.4% for +Var, 2.4% for +S.K, 6.2% for +Coch.Corr, 2.9% for +Mod.Power, 1.4% for +C1+C2, and 6.7% for Ori..

In the non-A1 proportions of multiunits with significant VE values were even smaller. The proportions for VE by type were: Power: 0, +Var: 4%, +S.K.: 1%, +Coch.Corr: 0%, + Mod.Power: 0%, +C1+C2: 0%, Ori: 3%. Those for VE by exemplar were: Power: 4%, +Var: 1%, +S.K.: 4%, +Coch.Corr: 6.9%, + Mod.Power: 5%, +C1+C2: 4%, Ori: 5%. These data thus confirm that the cortical responses we observed were generally very poor at discriminating texture types or exemplars types largely irrespective of the wealth of statistical features offered, in sharp contrast to the IC responses.

### Texture classification performance based on single-trial neural activity

The analysis so far has focused on the responses of individual multiunit, and has revealed that IC units are more sensitive to the statistical features that distinguish natural sound textures than AC units. Indeed, the ability of cortical units to code texture features or texture identity seemed surprisingly poor. However, perhaps the cortical encoding of texture features improves if neural population codes are considered. To examine this possibility, we asked how well single-trial neural ensemble responses can be classified according to texture type, and how the number of units in the neural ensemble and the duration over which responses in the ensemble are accumulated influence the classification performance. We might expect that the classification performance should increase as more units and longer time windows were included in the analysis, because the neural activity from more units can potentially encode more stimulus information, and accumulating responses over longer time windows may improve signal to noise ratios. While considering population codes may reveal more encoding of texture features at the level of the cortex than we had seen in the previous analysis of individual units, we would nevertheless expect IC ensemble classification performance to exceed that seen in the A1 and non-A1, given the widespread feature selectivity of IC neurons just described. To test these predictions, we trained a Gaussian decoder (see Methods) on the neural responses to the original sounds, and then quantified the classifier performance on single-trial neural responses to each texture morph. The classification performance (averaged over texture types) increased in the IC (Figure 5A), A1 (Figure 5B), and non-A1 (Figure 5C) as the number of multiunits and the analyzed time window increased (permutation test, p < 0.05). The classification performance based on IC neural responses was higher than in the A1 and non-A1 for all population sizes and durations considered in the analysis (permutation test, p < 0.05).

**Figure 5.**
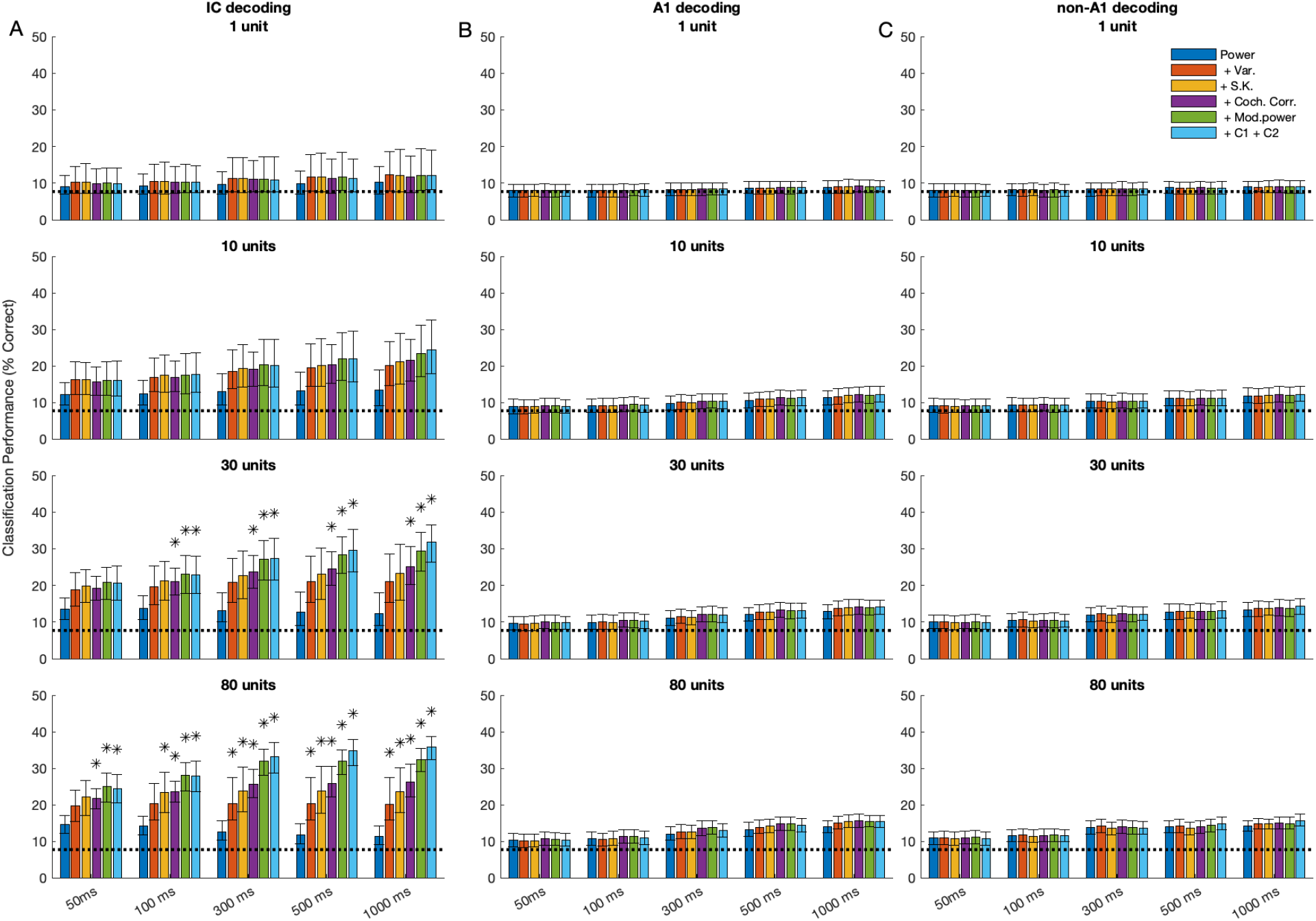
Mean classification performance over texture types as a function of the texture morphs, number of analyzed units, and the neuronal response time window. The classification performance in the IC (A), A1 (B), and non-A1 (C) increased with the time window and the number of units included in the analysis. The time window is indicated at the bottom of the figure, and the number of units is indicated at the top of each panel. Error bars represent the 2.5th and 97.5th percentile of the distribution of 1000 simulated ensembles. The dashed lines represent chance performance (1/13). Asterisks show conditions in which classifier performance was significantly higher than in the power condition (p < 0.05, permutation test). Different colors represent the classification performance for each texture morph (see legend in the upper right corner).

In the IC, the decoder showed significantly higher performance (permutation test, p < 0.05) for the neural responses to texture morphs including the higher-order statistics (+Mod.power and +C1 +C2 condition) than in the power condition, when subsets of 30 units and time windows longer than 500 ms (Figure 5A, the third row) were included in the analysis. Furthermore, the classification performance of IC units for adding more higher-order statistics in the texture morph (+Coch.Corr, + Mod.power, + C1+C2) was significantly higher (permutation test, p < 0.05) than in the Power condition, when subsets of 100 units and time windows longer than 50 ms were included in the analysis (Figure 5A, the fourth row). In contrast, for the A1 and non-A1 neural ensembles, the classification performance increased only very modestly or not at all (permutation test, p > 0.05) as texture morphs with higher order statistics were presented. The classification thus shows clear evidence for a high sensitivity to higher-order statistical features in the neural activity of the IC, but not in the auditory cortex.

The shortest response time windows for which classification performance above chance could be demonstrated with ensembles of 10 units was 50 ms in the IC but 500 ms in the cortex, respectively (Figure 5, the second row). In A1 and non-A1, this minimum classification time decreased to 100 ms when 80 units were included in the analysis (Figure 5B, the fourth row). In the IC, the minimum classification time stayed at 50 ms as more than 10 units were included in the analysis (Figure 5A, the second row). Longer integration times are therefore needed to achieve any degree of useful classification with ensembles of A1 and non-A1 units than with IC units, which may be at least in part due to the fact that cortical units typically have weaker sustained responses than IC neurons (compare examples in Figure 1).

## Discussion

In this study, we compared the way in which IC and AC encode statistical features of textures, and how that encoding may serve the purpose of texture discrimination. We found differences between IC and AC to be dramatic, in ways that may perhaps be surprising if one conceptualizes the auditory pathway as a processing hierarchy that proceeds from a simple encoding of acoustic features towards complex processing that facilitates the recognition of auditory objects. Overall, AC multiunits were not just a lot less likely to detect sudden changes in statistical features of sound textures, their responses were also less able to overtly discriminate between different texture types, or to generalize across different exemplars of a given texture type, than IC units. This may appear counter-intuitive if we make the assumptions, firstly, that ascending along a sensory pathway brings us closer to “where perceptual decisions are made”, and secondly, that the neural codes underpinning perceptual decisions are either simple rate codes or simple firing pattern codes that can be revealed at the level of firing of individual units (Parker & Newsome, 1998). Classic work (Salzman et al., 1990) on visual cortical streams would suggest that such assumptions are at least occasionally plausible, and even in the auditory cortex, there have been some successful attempts in decoding firing patterns of individual units based on the premise that greater perceptual dissimilarity of two complex sounds should go hand in hand with greater dissimilarity of neural firing patterns (Walker et al., 2008). When considered against these assumptions, our results may appear surprising, but when one casts a wider look at the relevant literature one notices that several recent studies are nevertheless in very good agreement with many of our key observations. We shall review some of these in the following paragraphs.

### IC responses are very good at encoding textures

In a previous study (Mishra et al., 2021), we had demonstrated that many IC neurons are sensitive to the changes of the statistical features of textures, and this sensitivity involves mechanisms beyond a simple linear dependence on the sound spectrogram change. In the current study, we were further able to show that many IC neurons encode more information about texture types than about individual exemplars, which suggests some level of generalization or invariance, which one might expect to see in “high-order” sensory neurons involved in auditory object classification rather than simply encoding acoustic stimulus parameters. We also found that the classification performance based on single-trial neural responses increased as more features were added to texture morphs (Figure 5A), and this trend is similar to what is observed in human psychophysical performance when human participants identify texture sounds (McDermott & Simoncelli, 2011). Our results suggest that IC neural responses would be well suited to informing texture recognition tasks, and this observation is consistent with a recent study by Zhai et al. (2020). In fact, statistics extracted from the textures were based on the auditory physiological properties of subcortical stages from the auditory periphery to the auditory thalamus, and the synthesized textures by these statistics are nevertheless perceptually similar to the natural textures (McDermott & Simoncelli, 2011). Thus, our findings that these statistics modulated the neural responses in the IC but not in the cortex support the notion that subcortical processing plays essential roles in texture recognition.

### AC responses are much less sensitive to texture features than IC neurons

In sharp contrast to the IC, AC responses typically showed only little or no sensitivity to changes in the statistical features of textures, did not discriminate texture types well, and were typically more sensitive to differences in texture exemplars than in texture types. Also, unlike for the IC, texture classification based on AC neural responses did not improve as more statistical features were added to the texture morphs. In that sense, our IC results are arguably more similar to human perception than our AC results, given the just mentioned trend for human texture identification performance to increase with the number of features present (McDermott & Simoncelli, 2011). In this respect, our AC results may appear surprising. They also contrast with recent studies of texture coding in the visual system: Ziemba et al. (2016) showed that responses of V1 neurons were highly informative in a visual texture exemplar classification, while V2 neurons showed better performance in texture type classification. In contrast, our A1 and non-A1 neurons were generally poor at texture type classification. But while our AC results may seem surprising when compared against IC or against visual cortex, they are in fact in good agreement with results reported in several other recent studies of cortical responses to complex, natural sounds. For example, our finding that the classification performance in AC was consistently worse than in IC is entirely consistent with a recent study by Souffi et al. (2020), which observed that neural discrimination performance across guinea pig vocalizations in guinea pig IC was higher than in guinea pig A1 or A2. Meanwhile, Blackwell et al. (2016) investigated responses of individual single units in the A1 of awake rats to synthesized, water-like texture stimuli, and found that many neurons appeared to be selective for only a subset of stimuli with particular spectro-temporal density and cyclo-temporal statistics. Their neurons thus also generalized poorly across different types of water sounds. Furthermore, DeWeese et al., (2003) noted that AC differs from visual cortex in that AC responses are generally much more transient and sparser. The example responses shown in Fig 1 of this study certainly suggest that A1 and non-A1 neural responses to textures are sparser than IC responses. Similarly, Montes-Lourido et al., (2021) studied the neural representations of different types of guinea pig vocalizations along the ascending auditory pathway, and found pronounced changes in neural responses as neural processing progressed from sub-cortical to cortical structures, which resembled the differences between IC and AC we document here. These authors observed that neurons in the ventral medial geniculate body and layer 4 of A1 were not very selective, in that they responded to most if not all of the vocalizations tested, and they fired throughout the call durations, while neurons in other, “downstream” layers of A1 L2/3 fired very sparsely, responding selectively to only very few vocalizations with short bursts of firing. Thus, the sparse cortical responses to textures that we see are not unexpected when compared against previous reports of cortical responses from other groups. Nor is it entirely surprising that cortical responses to textures appear less informative than IC responses, given that other previous studies have reported 2- to 4-fold decreases in stimulus related mutual information in AC compared to IC neural responses (Chechik et al., 2006; Schnupp, 2006).

### What might explain the weak responses and poor classification seen in cortex?

One question worth addressing is whether the poor texture classification exhibited by the cortical responses we recorded could be an artifact of anesthesia. It is of course difficult to generalize when discussing anesthesia effects given the large diversity of anesthetic agents available and the fact that effects are invariably dose dependent, and we do not doubt that anesthetics can play a big role, particularly if doses are high or agents are not well chosen. But it is also worth remembering that effects of anesthesia are not always detrimental to neural coding in the auditory pathway. For example, Ter-Mikaelian et al., (2007) found that anesthesia could enhance the temporal precision of A1 neuron responses, while having no clear effect on IC neuron temporal response properties. To us it seems highly unlikely that the poor texture discrimination we observed in the firing of cortical neurons is an artifact of the anesthesia we used, given that several of the studies just described (Blackwell et al 2016, Montes-Lourido et al 2021, Walker et al 2008) included data from awake, unanesthetised animals, and nevertheless observed cortical responses that were similar to those seen in in our anesthetised preparation. Walker et al (2008) even performed a direct comparison of anesthetised and unanestehtised data and found no significant differences.

What other reasons might explain the dramatic differences in texture coding observed between midbrain and cortex? Several authors have commented on the fact that AC neuron responses are often highly nonlinear, in ways that not only makes them less like V1 simple cells, but also sparser in their firing, with quite idiosyncratic response preferences (DeWeese et al 2003, Chechik and Nelken 2012, Bathellier et al. 2012). One potentially relevant finding here is that the majority of AC neurons appear to exhibit non-linearities that make them operate like logic AND gates (Harper et al, 2016), which would contribute to their idiosyncratic and sparse stimulus selectivity, but would not necessarily make them good at generalizing across exemplars in a stimulus class. Another is that AC neurons have a tendency to adapt strongly to ongoing “background” sounds (Rabinowitz et al, 2013, Robinson et al, 2016, Khalighinejad et al, 2019). This tendency could in some sense explain why cortical neurons seemed to be largely “uninterested” in the relatively long, stationary texture stimuli used in this experiment.

### What is the role of cortex in the perception of auditory textures?

Given that this study comprised no behavioral tasks and only collected and analyzed neural responses from anesthetized rodents, any interpretation of our data regarding how neural processing ultimately leads to perception of auditory textures must necessarily be quite speculative. On the surface of it, our results would seem to invite the conclusion that the auditory midbrain is much better placed to carry out pattern classification tasks than auditory cortex, given how much easier it is for us to decode IC responses to induce texture identity than AC responses. However, we must not jump to conclusions. Building classifier algorithms to analyze neural response patterns “externalizes” many key steps involved in auditory perception which the auditory brain would normally have to perform independently, such as consulting memory traces that form templates for recognizable texture types, or providing a behavioral context which dictates that perceiving, identifying or discriminating auditory textures is the task of the hour, and what to do with the outcome of this perceptual analysis. AC neurons may carry much less overt texture cue information in their responses, but they may be much better able to compare current stimuli against templates stored in memory in a context dependent manner. One previous study by Bathellier and colleagues (2012) argues that AC neurons likely encode stimulus identity through widely distributed population codes, and our own pattern classifier results are compatible with that idea. It is in principle possible that the pattern classification performance for our AC responses would improve somewhat if temporal pattern codes, rather than simple average rate codes, were analyzed, as has been successfully done in some previous studies (Narayan et al., 2006; Schumacher et al., 2011; Woolley et al., 2005; Walker et al 2008). However, a finely resolved temporal pattern analysis is bound to arrive in misleading results in the context of auditory texture analysis given that texture identity is characterized by stationary, time invariant statistics as opposed to precise temporal patterning: for example, we recognize applause as a stochastic grouping of claps, not a precisely timed rhythm of claps. We are therefore not pursuing temporal pattern codes here because doing so would bias the analysis towards identification of the specific patterns of particular exemplars rather than texture classes. Nevertheless, we must remain open to the idea that AC may have important roles to play in the perceptual processing of auditory textures, even if our results clearly show that individual units in AC display very little overt sensitivity to statistical texture features or to differences in texture types.

## Acknowledgments

This project was funded by the Hong Kong General Research Fund (No. 11100617). NH was supported by the Wellcome Trust (grant no. WT108369/2015/Z), and in accordance with the open access conditions of this grant, a CC BY public copyright licence is applied to any Author Accepted Manuscript. This work has been supported by the European Commission’s Marie Skłodowska-Curie Global Fellowship (750459 to R.A.) and a grant from European Community/Hong Kong Research Grants Council Joint Research Scheme (9051402 to R.A. and J.S.).

## Notes

### Competing Interest Statement

The authors have declared no competing interest.

